# Economical Method to Construct a Prism-based TIRF Setup on an Existing Confocal Microscope to Perform smFRET Experiments

**DOI:** 10.1101/2025.02.21.639412

**Authors:** Pratibha Agarwala, Dibyendu K. Sasmal

## Abstract

Total internal reflection fluorescence (TIRF) microscopy enables observation of complex bio-assemblies and macromolecular dynamics in high spatial-temporal resolution at single-molecule level in real-time. With TIRF illumination, fluorophores near a sample substrate are excited by an evanescent field, thereby circumventing the axial diffraction limit of light. Prism-based TIRF (p-TIRF) microscopes are comparatively easy to use. They may be readily adjusted to meet the requirements of a broad range of experimental applications, such as to examine of macromolecular complexes, to study of the behaviour of vesicles and small organelles, the study of protein-DNA complexes at the single-molecule level. These experiments can give unique insights into the mechanisms driving the molecular interactions that underline many fundamental activities within the cell by providing information on fluctuation distributions and unusual events. Here, we report a detailed method to build a p-TIRF setup inexpensively using an existing confocal microscope where the same light source can be used for both systems. Furthermore, we provide a brief overview that aims to give the readers a stepwise tutorial protocol for building, assembling, aligning, and preparing the specimen to conduct single-molecule fluorescence resonance energy transfer (smFRET) experiments using a custom-built p-TIRF setup. We believe that this article will be of assistance to labs that already have a confocal microscope and want to perform TIRF experiments for potential future applications.

**Research Highlight:** We report here a simple and cost-effective method to build a prism-based TIRF setup on a laser scanning confocal microscope. The setup is affordable, and users can use both confocal and TIRF modes in the same setup with the same light source and optical components.

**Graphical Abstract:** 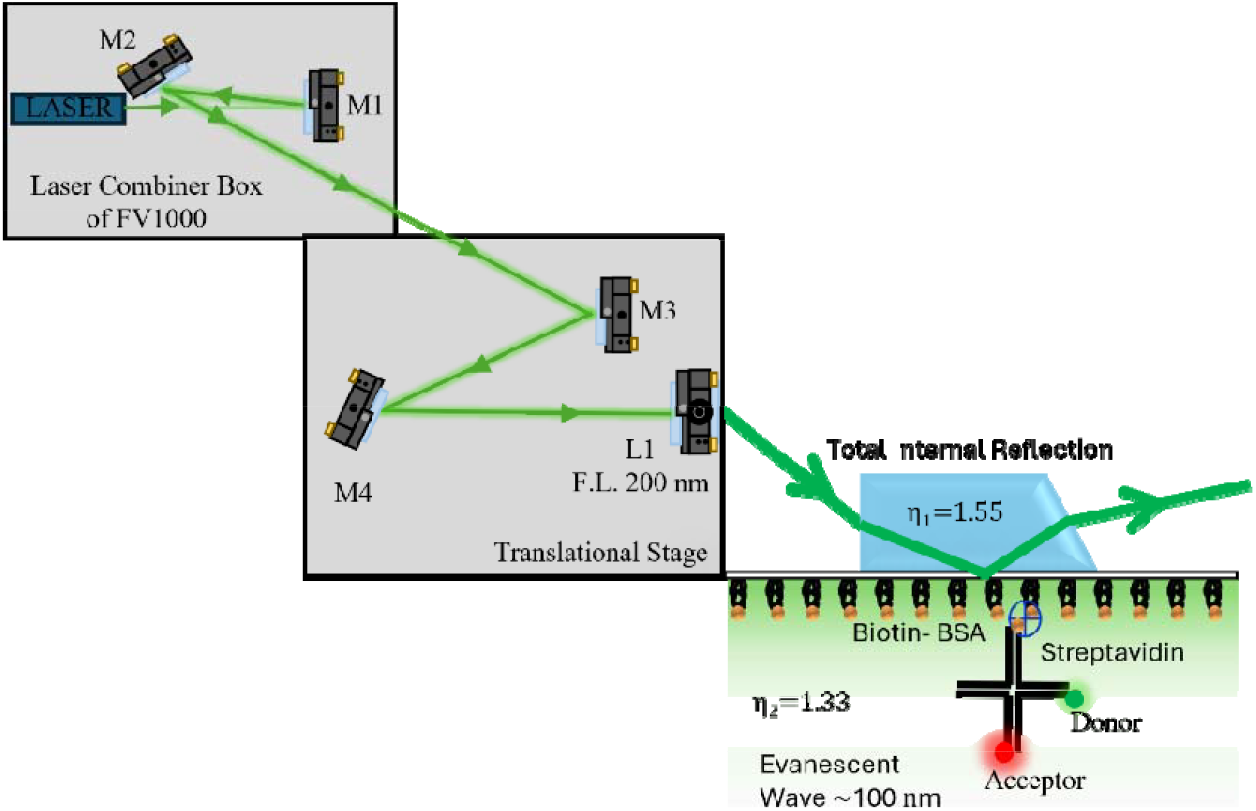

## Introduction

Total internal reflection fluorescence microscopy (TIRF) is a widely used imaging technique in the fields of cell biology and biochemistry to study single molecules.^1-5^ The concept of total internal reflection relies on an excitation light beam that travels to two different refractive index planes, and by the principle at a specific critical angle, the light is totally reflected from the glass/water surface, which creates an electromagnetic field.^6^ The electromagnetic wave has the same frequency as the incident light called the evanescence wave, which can penetrate a shorter distance of the sample by utilizing the phenomenon of total internal reflection to achieve high-resolution imaging of the molecule of interest.^5^ The depth of the evanescence field can reach approximately 100 nm, and it exponentially decreases as the penetration depth increases^6^, which depends on several factors, including the wavelength of light, the angle of incidence, and the difference in the refractive index of the medium.^5, 7^ Generating the evanescent wave has the advantage of minimizing photobleaching as the illumination volume decreases while also enabling the visualization of structures below the axial diffraction limit.^8^

Researchers widely use TIRF microscopy to observe cell contacts and investigate cell adhesion, ion channels, endocytosis, and filamentous protein self-assembly.^2, 9-10^ The total internal reflection microscope is also utilized to study the smFRET experiment, by which it is possible to precisely analyse the conformational change and molecular interaction of any biological system.^11-16^ There are two ways to develop a TIRF microscope: A) objective and B) prism-based total internal reflection microscopy. Although there are several advantages to the objective-based TIRF, e.g., the specimen here is fully accessible, and the user can easily switch between the epifluorescence mode and objective-based TIRF, which requires less complex alignment, it also has several disadvantages. Building an objective-based TIRF setup requires a higher numerical aperture objective, which can be more costly.^10, 17^ This objective allows confinement closer to the surface, as it focuses light at the back focal plane, so there is a possibility of light scattering within the objective, which reduces the signal-to-noise ratio.

The prism-based TIRF is the most sophisticated and cost-effective technique, providing distinct advantages, e.g., greater penetration depth as compared to objective-based TIRF.^18^ It also reduces the optical abbreviation that may arise from the objective lens, which improves the spatial resolution, enhances the signal-to-noise ratio, and provides us with greater experimental flexibility by fine-tuning the excitation beam. In the prism-based TIRF, the field of illumination is higher as compared to the objective-based TIRF.^4^ Here, we use a prism to attain the critical angle, directing the laser light between the glass and water media at a specific incident angle that can cause total internal reflection.^10^

Despite the numerous applications, there are currently no commercial instruments available for the prism-based TIRF microscope, which demands a substantial amount of skill to operate such sophisticated equipment.^17, 19-20^ Here, we directly address a step-by-step guide to building the p-TIFR instrument using a confocal microscope. This study gives an overview of the method used to create p-TIRF systems.

## Materials & Methods

**Table 1:** Extensive list of microscope parts and chemicals.

| Item Description | Catalogue No | Vendor |
| --- | --- | --- |
| Standard Solid Breadboards | BB-300400-AL | Holmarc India |
| 12 mm Diameter Posts | P-75, P-100, P-250 | Holmarc India |
| Post Holders for 12 mm diameter Posts | PH-75, PH-100, PH-250 | Holmarc India |
| H Type Post Base | PB-7525 | Holmarc India |
| Square Type Post Base | PB-6550 | Holmarc India |
| Table Clamps (L-shaped) 50x15x16 mm | L-TC-50 | Holmarc India |
| Plano Silver Mirror | HO-PCMS-25 | Holmarc India |
| Biconvex lens | HO-BXL25-08B | Holmarc India |
| Pellin Broca Prism<br>Clear aperture = 5 mm , Material = UV<br>Fused silica | HO-PBS-5 | Holmarc India |
| Kinematic Mount for Circular Optics | KMC-CS-25 | Holmarc India |
| Translation Stage | TSQ90-Mu5-03 | Holmarc India |
| Swivel post clamp | SWPC12 | Holmarc India |
| Right angle post clamp | RAC-12 | Holmarc India |
| Rigid Flip Mount | FLRMM-25 | Holmarc India |
| Right angle Post clamp | RAC-12 | Holmarc India |
| Post Collar | PC-12 | Holmarc India |
| Metric Socket Cap Screws | M4SHC16 | Holmarc India |
| Metric Socket Cap Screws | M6SHC16 | Holmarc India |
| Metric Set Screws | M4SCGB16 | Holmarc India |
| Metric Set Screws | M6SCGB16 | Holmarc India |
| Honeycomb Tabletop with vertical and<br>horizontal vibration isolation |  | Holmarc India |
| Quartz Slide | 42297 | Alfa-Aesar |
| Double-sided tape Thickness (100 μM) | 93010 | 3M |
| Immersion oil | 56822 | Sigma Aldrich |
| 5-minute Epoxy glue |  | Local Vendor |
| Biotin BSA | 9048-46-8 | Sigma Aldrich |
| Streptavidin | 434301-1MG | Thermofisher |
| Trolox | 238813-1G | Sigma Aldrich |
| Glucose oxidase | 9001-37-0 | Sigma Aldrich |
| Catalase | 9001-05-02 | Sigma Aldrich |
| ssDNA |  | Sigma Aldrich |
| Laser Scanning Confocal Microscope |  | Olympus FV1000 |
| EMCCD Camera with Optosplit - II |  | Andor iXON 897, 2021 |
| Bandpass Filter | ET585/65m, ET706/95m | Chroma Technology |
| Dichroic Beamsplitters | ZT633rdc-UF2 | Chroma Technology |
| Diamond drill bit 1.0 mm |  | Amazon |
| Black Mounting board 1mm thickness |  | Amazon |
| Electric Drilling Machine |  | Amazon |
| Tubing | 0.6 mm I.D. x 1 mm O.D. | Local Vendor |

### Microscopy Setup

We developed a custom-made prism-type TIRF (p-TIRF) setup on top of an Olympus Confocal Microscope (FV1000). First, we installed an anti-vibrational optical table with an overhead shelf to minimize table vibration and ensure proper alignment. We also chose a dark and dust-free room to minimize the noise from the stray light and the amount of dust that accumulates over time. We extracted the laser from the FV1000’s laser combiner (Figure 1A). The laser combiner box, which connects to the scanner head via fibre optics during confocal imaging, consists of six lasers. We positioned a flip mirror M1 in front of the final mirror before the fibre optics, ensuring that the laser power was at its peak. To prevent dust particles, we attached black cardboard to the side of the laser combiner box, and we cut one side of the cardboard to allow the flip mirror to access it. After the laser beam reflected through the flip mirror, it was pointed towards mirror M2, which was positioned vertically such that the beam could travel in the direction of the breadboard where M3 was located (Figure 1B). The laser beam then proceeded to an elevated breadboard situated 30 cm above the optical table.

**Figure 1.**
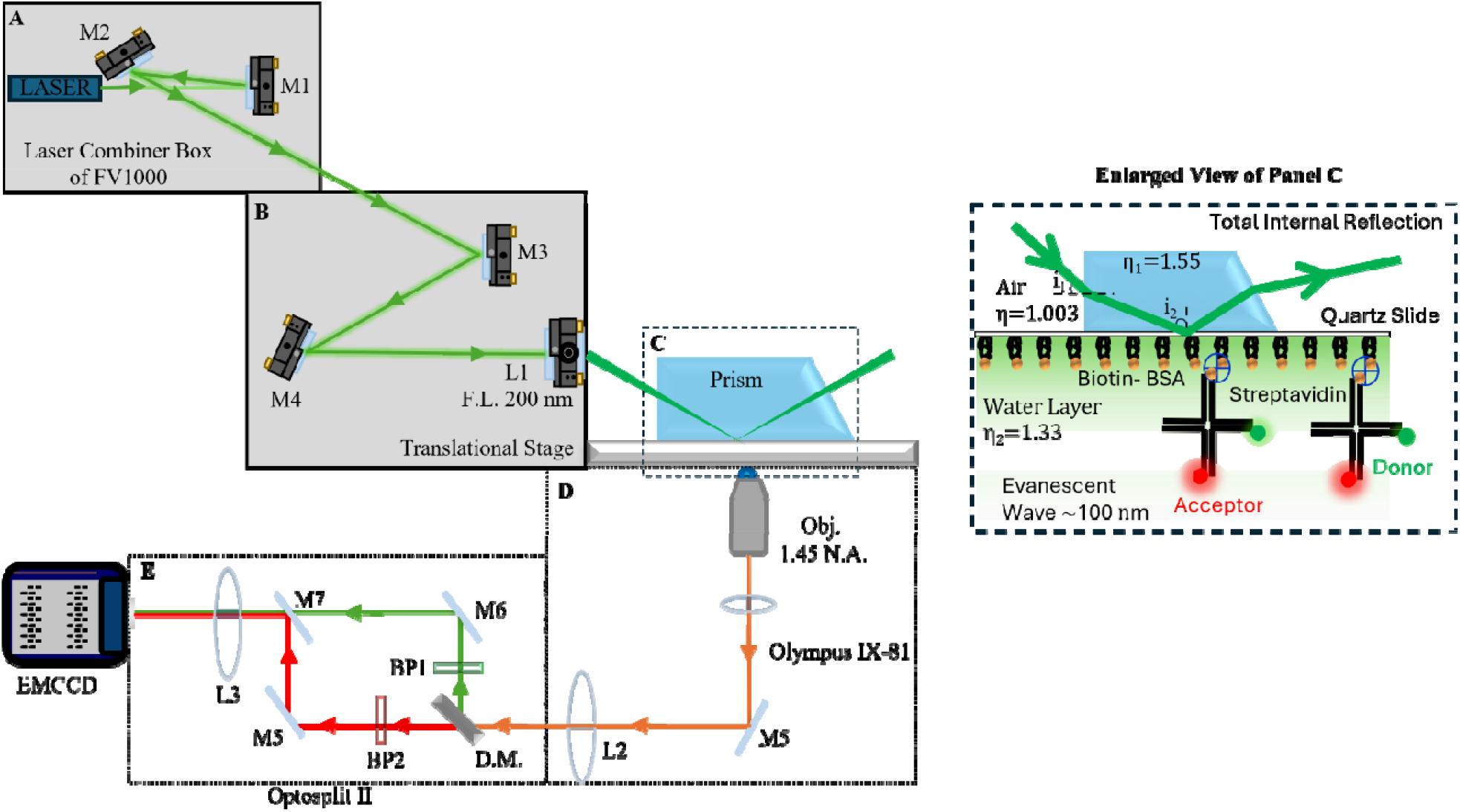
Layout of prism-based TIRF microscope. The complete system is divided into five sections: **(A-B)** the excitation part from the laser combiner box and at the elevated breadboard for focusing the laser beam, respectively. **(C)** The prism is attached to a prism holder mounted with a prism holder carried arm (discussed in Figure 2). In the enlarged view, we have shown the light path at the interface of the quartz/water surface, which creates almost ∼100 nm evanescent wave to excite the fluorophores. **(D)** The emission path: The fluorescence signals from the sample are collected by a 100X oil-immersion objective with 1.45 N.A. and pass towards th optosplit. **(E)** Schematic representation of optosplit II. Dichroic Mirror and emission filters were used to separate the emission channel of cy3 and cy5 in optosplit II and projected onto th EMCCD camera.

Installation of the XYZ translation stage on the elevated breadboard takes place on the left side of the microscope and top of the optical table (Figure 1B). A distance that enables the translational stage to preserve the focusing lens’s focusing distance should be using elevates the breadboard parallel to the microscope stage height. We secured the optical table with an H-type clamp to ensure stability. A custom-built prism holder carried arm, discussed in the next section, mounts the prism over the microscope stage. Precise changes to focus the laser above the objective are made possible by the 1 cm x, y, or z movement of the XYZ translational stage, to which the focusing lens was attached. To focus the laser more precisely, a plano-convex mirror with a 200 mm focal length, mounted in an XYZ translation stage, fine-tuned the laser beam to create the total internal reflection.

To position the lens, we first insert it into a lens mount (Figure 1B), which also features a fine-tuning knob and a post. Then, during the translation stage, we insert it into the post holder, ensuring that the flat side faces the objective. Briefly, after receiving the laser beam from the flip mirror, the second mirror leads the laser towards the elevated platform (breadboard). We fixed two additional mirrors so that the final mirror on the breadboard focused the laser through the 200 mm lens towards the prism. We position the next two mirrors on the breadboard at approximately a 45-degree angle to focus the laser beam onto a 200-mm focal length lens. The laser path should enter the Pellin-Broca prism at the appropriate angle so that it can create the total internal reflection. We used a prism to create the total internal reflection. As a result, the incident beam can be fully internally reflected, creating an evanescent field on the quartz-water(sample) interface (Figure 1C). It was essential to keep in mind to use a non-fluorescent immersion oil between the quartz slide and the prism that has the same refractive index as the slide.

For our setup, the angle of the laser beam falling on the prism was approximately 31°. At this point, it was important to control the field of view of the excitation area by placing the lens in the correct position. For the laser alignment, first focus the 0.1 µm fluorescence bead on the flow chamber (discussed in the next section) to fix the objective height using the epifluorescence light. Once you can see the sample through the eyepiece, proceed to align the laser. Next, aim the laser at the top of the objective, ensuring it is roughly focused. After this, apply a small drop of non-immersion oil to the top of the sample chamber and position a prism with a prism holder carried arm on top of it.

To achieve this, we secure the first prism holder carried arm on the microscope body and loosen the prism holder’s (the technical details are described in Figure 2) screw to adjust the prism position during the alignment process easily. Next, we place the prism on top of the sample chamber, ensuring that it remains flat against the sample chamber. After that, we performed a rough alignment to reflect the laser beam from the prism, ensuring that the field of view was centred in the lower objective through the eyepiece. With the completion of the alignment, we tighten the prism holder screw to secure the prism with the adapter. Fixing the prism to the microscope body was crucial to prevent the excitation beam from moving when the sample moved in the x-y direction to collect images in different areas. Following this, we primarily utilize the transitional stage for fine alignment each time we swap out the sample. Every time we experiment with a new sample chamber, we must first remove the prism adapter to position the sample chamber above the objective, necessitating a fine alignment. To capture the image, we currently use the EMCCD camera and the micromanager software. Next, we utilize the translation stage to align the Z-axis, ensuring the bead sample reaches its maximum intensity. We also align the x and y axes to maintain the centre of the field of view.

**Figure 2.**
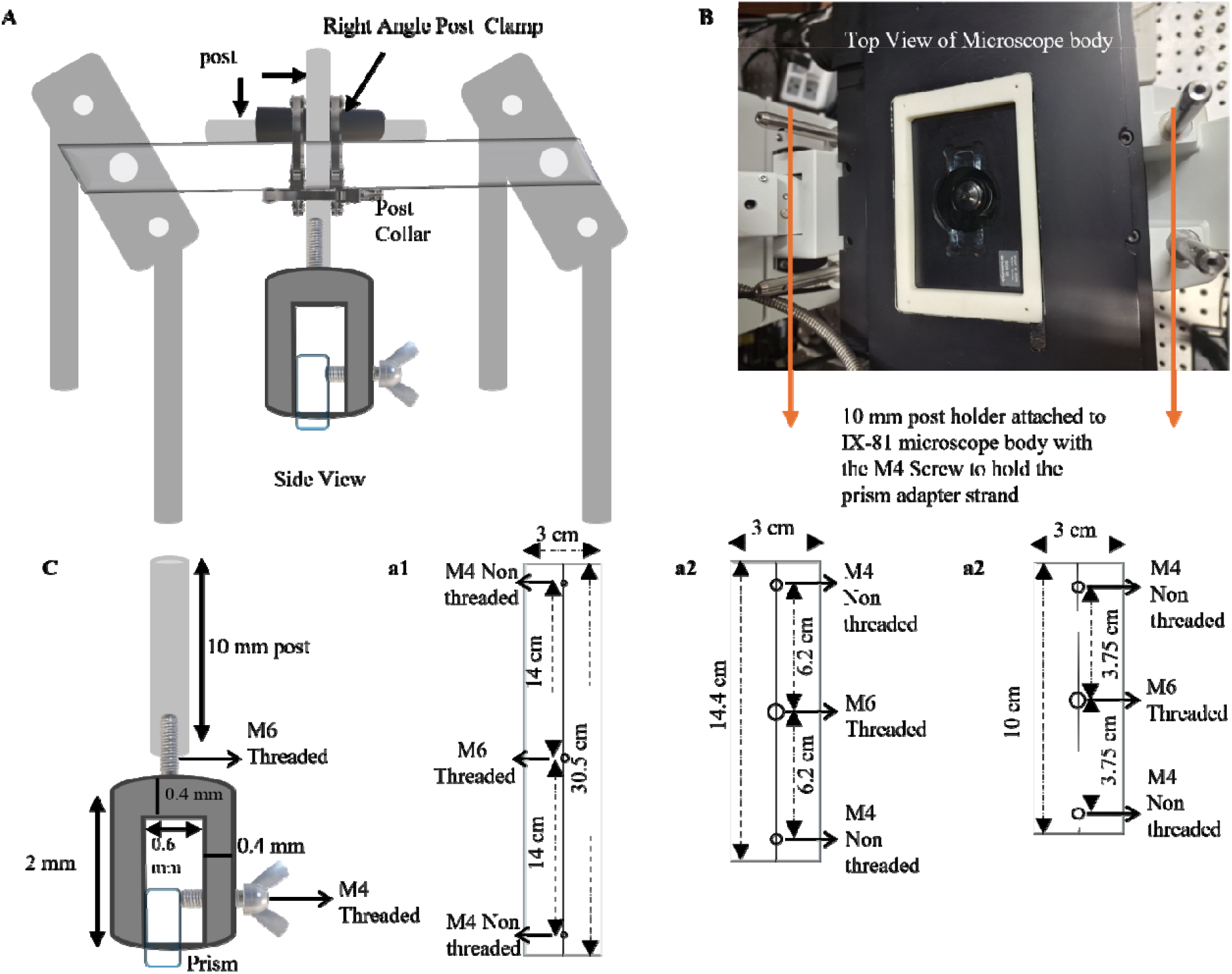
Schematic representation of prism holder and prism holder carried arm. **(A)** The dimension of the prism holder carried arm which consist of 3 aluminium plates as shown in a1, a2 and a3. **(B)** Real image of the top view of the microscope body used to hold the prism holder carried arm **(C)** The dimension and layout of the prism holder.

### Calculation of Critical Angle and the depth of penetration

We have also calculated the incident laser beam angle for the total internal reflection. According to Snell’s law^7-8, 17, 19^

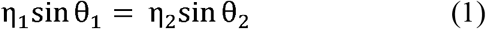

Where the incident beam and refracted beam are represented by θ_1_ and θ_2_ respectively, at the glass/water interface, and the refractive indices of medium 1 (glass surface= 1.55) and medium 2 (water = 1.33) are represented as η_1_ and η_2_, respectively. The quartz/water surface’s critical angle is

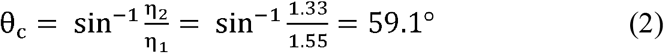

Therefore, in accordance with Snell’s law, the i_2_, as shown in Figure C, must equal or exceed 59.1 in order to produce a total internal reflection. For our setup, the angle of the laser beam falling on the prism was i_1_∼31°. So, as per the calculation, the i_2_ was approximately 71°, which was greater than the critical angle of the glass/water surface. We have also calculated the depth of penetration using the equation.^8, 19^

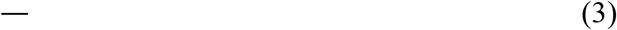

As the is the incident light’s wavelength (for our case, 559 nm), the depth of penetration was approximately 73 nm. However, the incident angle and the medium’s refractive index have a significant impact on the penetration depth.

### Prism Holder and Prism holder carried arm

A holding device (Figure 2c) with a prism-carrying arm (Figure 2A) mounted to the top part of the microscope was used to secure the prism above the flow cell. We have used aluminium machining to create our custom prism holder and holder carried arm. The holder carried the arm’s measurements, and the prism’s height will vary depending on the microscope. Still, the design’s primary idea—fixing the prism’s location directly over the objective remains constant. We internally created our prism holder and prism holder carried arm, as shown in Figure 2. The prism we tightened the Prism (Pellin-Broca, 11 mm x 20 mm x 6.4 mm) to the holder using M4 screws, keeping in mind that the holder size only accommodates a specific prism; for other prism sizes, we need to modify the design slightly. We have mounted the prism holder onto the holder carried arm with a post as per Figure 2 and used a right-angle post clamp and post collar to fix it. Although the prism holder and prism holder carried arm designed for our setup, readers may utilize it as per their setup. In our scenario, we affixed the 20 mm post holder to the IX81 microscope body using an M4 screw, followed by the attachment of the prism holder carried arm; (Figure 2 details all the dimensions. We determine the precise location for the screw head to fix the holder based on the dimensions of the setup and the location of the prism that we must mount to achieve TIR.

### Emission path

We collected the fluorescence emission using the confocal microscope’s transmission mode. We collected and separated the emission data from the excited fluorophore using the Optosplit II. First, we collected the emission from the 1.45 N.A. 100X objective to a tube lens (1X) in the FV1000, and then used the mirror (M5) to reflect it towards the optosplit. A C-mount adapter connects the optosplit to the left port of the FV1000. A filter cube made up of the dichroic mirror D.M. (chroma: ZT633rdc-UF2) is inserted by the optosplit. This separates the donor and acceptor signals before they go through the bandpass filters (B.P.): ET585/65m for the donor channels and ET706/95m for the acceptor channels. The focal lengths of L2 and L3 were the same, creating a 4F relay system for clear transmission of the microscope image. We connect the optosplit electron-multiplying charge-coupled device to the exit port and mount it on the optical table. The data collection software allows us to rotate the camera image either anticlockwise or clockwise. We primarily used Micromanager^21^ software to record the video in .tiff format and then processed the data using SPARTAN software.^22^ The manufacturer may provide detailed information about the camera manual or the data processing and collection software manual.

### Sample chamber preparation

We made sample chambers using a sandwich assembly of quartz slide and cover glass and sealed them with double-sided tape. The procedure is described below. ^23^

1. Take a quartz slide (76.2 × 25.4 × 1.00 mm) and create two diagonal holes by using the diamond drill bit of 1 mm.
2. After creating the hole, thoroughly clean the slide as described in the cleaning procedure.
3. Cut the double-sided tape diagonally (of approximately ∼ 5 mm widths) in the middle, as described in Figure 3.
4. Place one side of the double-sided tape on the slide and cover the other side with the coverslip to create a sample chamber of around 30-40 µl. To seal the chamber and remove any air bubbles properly, gently press the coverslip.
5. Cut 200 µl tips that are approximately 1 mm long. Next, place 200 µl of the pipette tip into each of the quartz slide’s two holes. This establishes a route for the exchange of solutions at the inlet and output. Finally, seal the parts with 5 minutes of epoxy glue.
6. On the inlet side, use a pipette to insert the sample, and on the other side, attach a tube of (0.6 mm I.D. x 1 mm O.D.) to create suction with a 1 ml syringe.
7. This way, you can modify your sample as needed.
8. Caution! Carefully inject the sample to ensure that no air bubbles enter the chamber.
9. After using the sample chamber, place it in water for at least 12 hours to remove the double-sided tape, and you can reuse the quartz slide.

**Figure 3.**
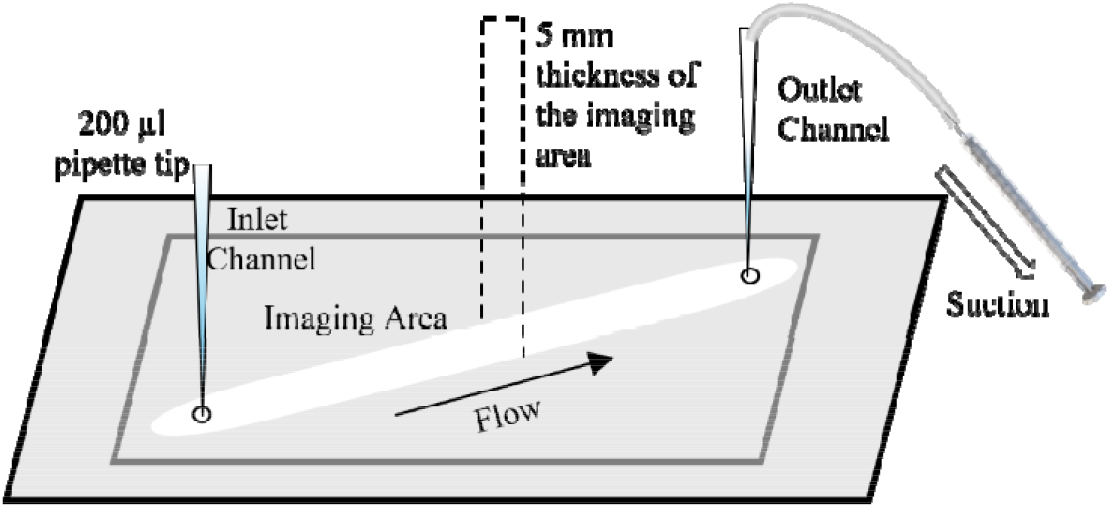
Schematic representation of sample chamber. The two corners of the quartz slide were drilled with the help of a diamond drill bit. Using double-sided tape (thickness 100 µM) as a spacer, a sandwich construction comprising the cover slide and cover glass was used to build the sample chamber. The two sides hole was then attached with a 200 µl pipette tip using the help of 5 minutes epoxy resin. A pipette was used to flow the channel, and a 1 ml syringe was used to create suction in the outlet area. The arrow shows the flow direction of the sample.

### Cleaning Procedure^24^

1. Boil the slide in MiliQ water for approximately 10 minutes.
2. Sonicate the slide in 0.1N NaOH for 15 minutes, NaOH is important to remove any biological residue.
3. Wash with tap water by gently scrubbing the slide.
4. For 15 minutes, sonicate the slide in acetone to get rid of any organic residue, followed by washing with Milli Q water.

### KOH etching Procedure

1. The slide and coverslip were both sonicated in 1M KOH solution for 20 minutes.
2. The slide and coverslip were both rinsed and sonicated with MiliQ water for 15 minutes.
3. Sonicate 20-30 minutes again the slide in 1M KOH and wash with water to make the surface hydrophilic by increasing the negative charge on the surface.
4. Dry the coverslip and slide with N_2_ gas to avoid any water spots in the slide, and burn the slide with a propane torch for 2 minutes to burn any leftover fluorescent residue.
5. Burn the coverslip with a propane torch for 20-30 seconds only; it may break if it burns for too long.

### DNA Holiday Junction Preparation

We purchased all the modified and unmodified ssDNA from Sigma Aldrich (custom-made) and used it without further purification. The sequence of four single-strand DNAs is given as follows. Cy5 labelled: 5′-Cy5-CCC TAG CAA GCC GCT GCT ACG G-3′, Cy3 labelled: 5′-Cy3-CCG TAG CAG CGC GAG CGG TGG G-3′, Biotin labelled: 5′-Biot-CCC ACC GCT CGG CTC AAC TGG G-3′, Unlabelled: 5′-CCC AGT TGA GCG CTT GCT AGG G-3′. We prepared the DNA Holiday junction by thermally annealing the ssDNA at a concentration of 2.5 µM in annealing buffer (10 mM tris HCl at pH=8.0, 50 mM NaCl, and 15 mM MgCl_2_,6H_2_O).^25^ We determined the cy3:cy5:biotin: unlabeled ratio to be 1:2:1:2. We carried out the annealing process by lowering the solution’s temperature from 96°C to 4°C at 0.5°C /min in a thermal cycler. We stored the assembled junction at 4°C for future use.

### Preparation of Oxygen Scavenger Solution (Trolox) ^23^

Approximately 30 mg of Trolox powder was dissolved in 5 ml of water using vortex; we were able to create the stock solution of (∼4 mM) Trolox. To achieve better solubility, we added 1 M NaOH drop by drop to the solution. After mixing, to estimate the exact concentration, we used a 0.2-μm syringe filter to filter the solution. Then, we examined the absorption spectra at 290 nm (with an extinction coefficient of 2350 ± 100 M^−1^ cm^−1^). Trolox is utilized as a triplet-state quencher to increase the dye’s photostability.

### Preparation of Imaging Buffer (T50) Solution^26^

T50 buffer is a mixture of 10 mM Tris-HCl and 50 mM NaCl at pH 8.0. For the Imaging Buffer preparation to avoid any impurities, we prepared 0.2 mg/ml stock catalase solution in T50 buffer. Based on this, we combined 10 mg of glucose oxidase with 20 µl of 0.2 mg/ml catalase in 80 µl of T50 buffer. In order to mix the ingredients, we centrifuge at a higher speed for one minute, extract the supernatant, and store it at 4 ºC. In this way, 100X gloxy solution was prepared. We prepared 0.8 % w/v D-glucose in the T50 buffer. We added Trolox and gloxy solution in such a way that the final solution of the imaging buffer contains 2 mM of Trolox solution and 1X gloxy. Careful! Just before the measurement, with the minimal amount of air exposure possible, we must add glucose to the glucose-containing buffer.

### Immobilization techniques^25, 27^

To immobilize the biotinylated DNA HJ to the quartz slide, 30 µl of Biotin BSA (1 mg/ml) was pipetted into the sample chamber. It was suctioned to the chamber with the help of a syringe. After 5 minutes, it was flushed with 100 µl of T50 buffer at least three times. Then again, using the same procedure, 0.2 mg/ml streptavidin was introduced, and after 1 minute, it was flushed with 100 µl of T50 three times to ensure the removal of any nonspecific binding.

After that, we added biotinylated DNA, waited for 3 minutes, and the sample was flushed with 200 µl imaging buffer to remove any unbound molecules. We generally used 100-200 pM DNA HJ solution for our experiment, although it highly depends on the setup, the optimal number of molecules, and other things. After the laser has been aligned with a standard bead sample, adjust the field of view by positioning the sample chamber on the stage adapter.

### Experimental Result

We have carried out a smFRET experiment to monitor the dynamic pattern of the Holliday junction (H.J.) to verify our experimental setup. The formation of the Holliday junction, a four-junction that is crucial to the repair of DNA breaks, occurred during homologous recombination.^28-29^ We have prepared the Holliday junction from the four ssDNA as described above. It was observed that in the presence of a divalent cation, H.J. shows conformational changes as the negatively charged phosphate backbone is charge neutralized by the divalent salt and electrostatic repulsion is reduced. Figure 4 reveals the representative trajectory of the DNA HJ conformational changes, which shows that it has a transition from a high FRET to a low FRET state in the presence of 50 mM Mg^2+^ concentration. The anti-correlation of the donor (green) and acceptor (red) trajectories was clear proof of a single molecule. ^30-31^ The data was processed in Spartan 3.7 software,^22^ and anticorrelated trajectories of the donor and acceptor were used for further analysis. The formula is used to further calculate FRET efficiency.^18^

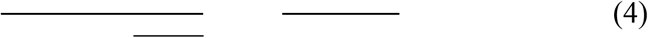

where I_A_(t) and I_D_(t) are the donor and acceptor intensities, and is the FRET efficiency. Here, the respective emission quantum yield of the donor and acceptor is represented by and Φ_A_ and donor and acceptor detector efficiency are symbolized by η_D_ and η_A_. We have considered that the correlation factor is 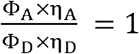.

**Figure 4.**
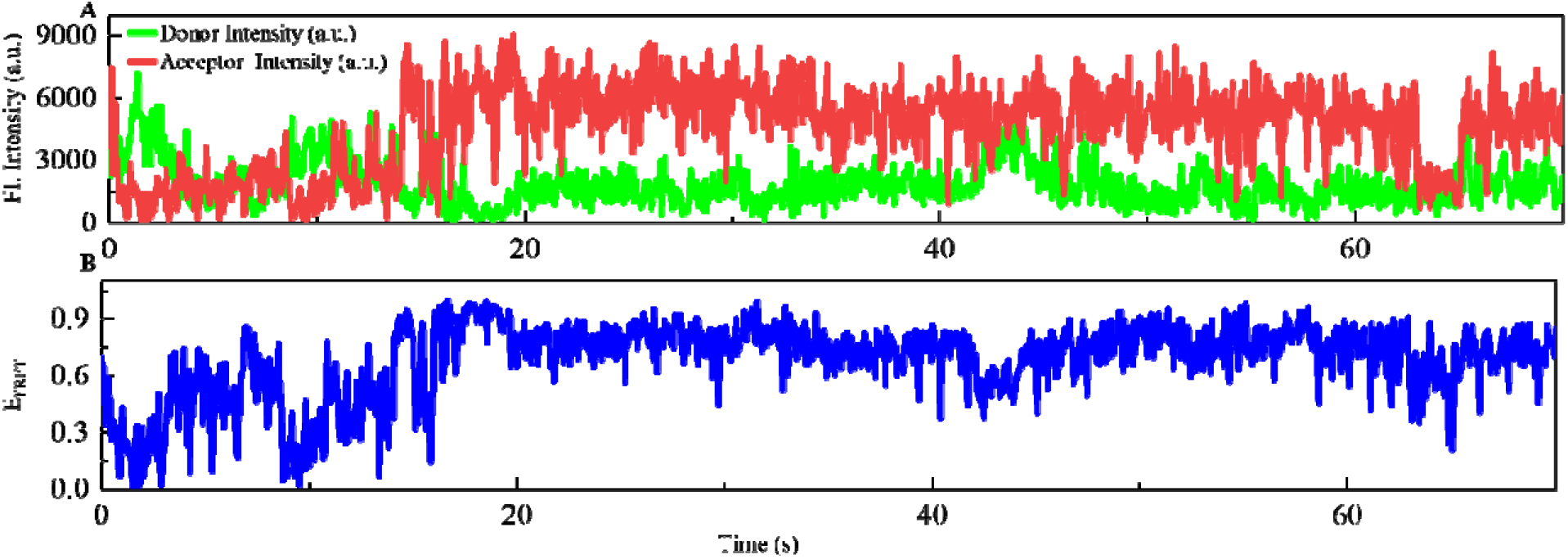
Validation of prism-based TIRF microscope by the smFRET experiment. **(A)** Representative plot of single molecule time trajectory was shown conformation switching of H.J. labelled with Cy3 and Cy5. **(B)** FRET efficiency fluctuation (blue line) calculated from the data shown in (A).

## Conclusion

Here, we have detailed a cost-effective protocol for constructing a p-TIRF microscope using a confocal microscope where both the confocal and TIRF microscope can use the same light source. We have also described in detail the preparation procedure for buffer samples, which can reduce photobleaching and photo blinking, as well as how to prepare the flow chamber. In our experimental results, we have seen a conformational change of DNA Holliday junction, which reveals that the system can perform smFRET to study the structural dynamic of biological samples.

## AUTHOR INFORMATION

### Corresponding Author

**Dibyendu K. Sasmal -** Department of Chemistry, Indian Institute of Technology Jodhpur, Email –, Phone: (+91)-291-280-1314, Web: www.sasmallab.org

### Note

Authors declare no competing financial interest.

## ACKNOWLEDGMENT

DKS thanks ANRF, Department of Science and Technology for Start-up Research Grant (Grant number – SRG/2020/001730) and Ministry of Education for STARS Grant (STARS2/2023-0473). DKS would also like to thank IIT Jodhpur for SEED Grant. DKS thanks Dr. Rajeev Yadav (Michigan State University) for valuable suggestions to develop pTIRF microscope. PA thank to IIT Jodhpur for fellowship. The authors also sincerely acknowledge help from Prof. Santanu Bhattacharyya (Former Director, IITJ) and Ann Wheeler (University of Edinburgh) for help and support to develop the microscope setup.

## Notes

### Competing Interest Statement

The authors have declared no competing interest.

